# A machine learning toolbox for the analysis of sharp-wave ripples reveal common features across species

**DOI:** 10.1101/2023.07.02.547382

**Authors:** Andrea Navas-Olive, Adrian Rubio, Saman Abbaspoor, Kari L. Hoffman, Liset M de la Prida

## Abstract

The study of sharp-wave ripples (SWRs) has advanced our understanding of memory function, and their alteration in neurological conditions such as epilepsy and Alzheimer’s disease is considered a biomarker of dysfunction. SWRs exhibit diverse waveforms and properties that cannot be fully characterized by spectral methods alone. Here, we describe a toolbox of machine learning (ML) models for automatic detection and analysis of SWRs. The ML architectures, which resulted from a crowdsourced hackathon, are able to capture a wealth of SWR features recorded in the dorsal hippocampus of mice. When applied to data from the macaque hippocampus, these models were able to generalize detection and revealed shared SWR properties across species. We hereby provide a user-friendly open-source toolbox for model use and extension, which can help to accelerate and standardize SWR research, lowering the threshold for its adoption in biomedical applications.

## Introduction

The study of brain rhythms has bolstered our understanding of the neural basis of cognition. Because these signals emerge from the coordinated activity of multiple neurons, they can be used as biomarkers of the underlying cognitive process^1^. For example, hippocampal sharp-wave ripples (SWRs) represent the most synchronous pattern in the mammalian brain, and are widely considered to contribute to the consolidation of memories^2^. SWRs consist of brief high-frequency oscillations or ‘ripples’ (100-250Hz), which can be detected around the hippocampal CA1 cell layer during rest or sleep. An avalanche of excitatory inputs from the CA3 region, typically visible as a slower sharp-wave component, triggers ripples locally in CA1^3,4^. Within the ripple event, neural firing patterns that occurred during exploratory behavior are reactivated outside of the experience^5,6^, leading the SWR to be used as an index of consolidation-associated reactivation or replay^7–10^.

Although SWRs can be detected across an array of recording methods, subfield locations, and species^2,11^ their underlying mechanisms and consequent local field potential (LFP) features are understood almost exclusively from measurements in rat and mouse dorsal hippocampal CA1. Even within this region, SWRs exhibit a large diversity of waveforms that presumably reflect the myriad combinations of reactivating ensembles^12–14^. Using spectral methods their characteristics are shown to vary along the long (septotemporal) CA1 axis within animals^15^ and most notably with phylogenetic distance across species e.g. when measured in the human versus non-human primates^11,16,17^. Furthermore, in diseases affecting hippocampal function, such as in Temporal Lobe Epilepsy, pathological forms of ripples have been reported ^18–21^, as well as along aging ^22,23^. However, spectral properties alone are suboptimal to separate these events from other types of faster oscillations ^24–26^.

To address this challenge, many researchers have developed feature-based strategies for detecting LFP oscillations using machine learning (ML) tools^16,27–32^. These novel strategies have accelerated our understanding the underlying mechanisms of SWRs, and the improvement of closed-loop interventions beyond those using spectral features alone^31,33^. Yet these methods have been focused on a single detection method optimized for a single target application, typically either in mouse dorsal CA1 or within lab-specific approaches to detection in brains of humans with epilepsy. As LFP recordings are increasingly common in the clinic, the need to scale analysis from small laboratory animals to the human brain is pressing^10,34–39^. Developing these new tools will provide the community with straightforward methods to identify SWRs from pathological oscillations across the range of recording technologies, sampled regions, and background pathologies. Therefore, there is a broad demand for a consolidated toolbox of ML methods for LFP feature analysis that can be easily applied across species, to aid in understanding of brain function, but also advance biomedical applications.

Here, we develop and analyze a set of ML architectures applied to the problem of SWR identification, and compiled in an open toolbox: https://github.com/PridaLab/rippl-AI. To favor an unbiased screening of potential ML solutions, we ran a hackathon with people from very disparate fields with the mission of detecting SWR using algorithms in a supervised manner. Using community-based solutions in neuroscience is gaining traction due to their ability to foster interdisciplinary and diverse perspectives, and to promote collaboration and data sharing^40–43^. We selected the most promising architectures from the hackathon and standardized them for fair comparisons. We show how the different ML models could bias SWR detection and identify conditions for their optimal performance and stability in the mouose hippocampus (*Mus musculus*). We then extend the analysis to SWRs recorded in the macaque hippocampus (*Macaca mulatta*), to demonstrate the generalizability of SWRs detection methods to the primate order. This proof of principle will foster the development of feature-based detection algorithms for future applications to a range of models and approaches, including the human brain.

## Results

### Community-based proposal of ML models of SWR

To create a diversity of ML supervised models of SWRs, we organized a hackathon that promoted unbiased community-based solutions from scientists unfamiliar with neuroscience research, and SWRs in particular (see Methods). The hackathon challenge was to propose a ML model that successfully identifies SWR in a dataset of high-density LFP recordings from the CA1 dorsal hippocampus of mice, used before for similar purposes^31^. Preparatory courses introduced participants into the main topics required for the challenge (Fig. 1A). To standardize the different ML models, they were given access to Python functions for loading the data, to evaluate model performance, and to write results in a common format. Annotated data consisted of raw LFP signals (8-channels) sampled at 30 kHz, and containing SWR events manually tagged by an expert (training set: 1794 events, two sessions from 2 mice; test set: 1275 events; two sessions from 2 mice; Fig. 1B).

**Figure 1:**
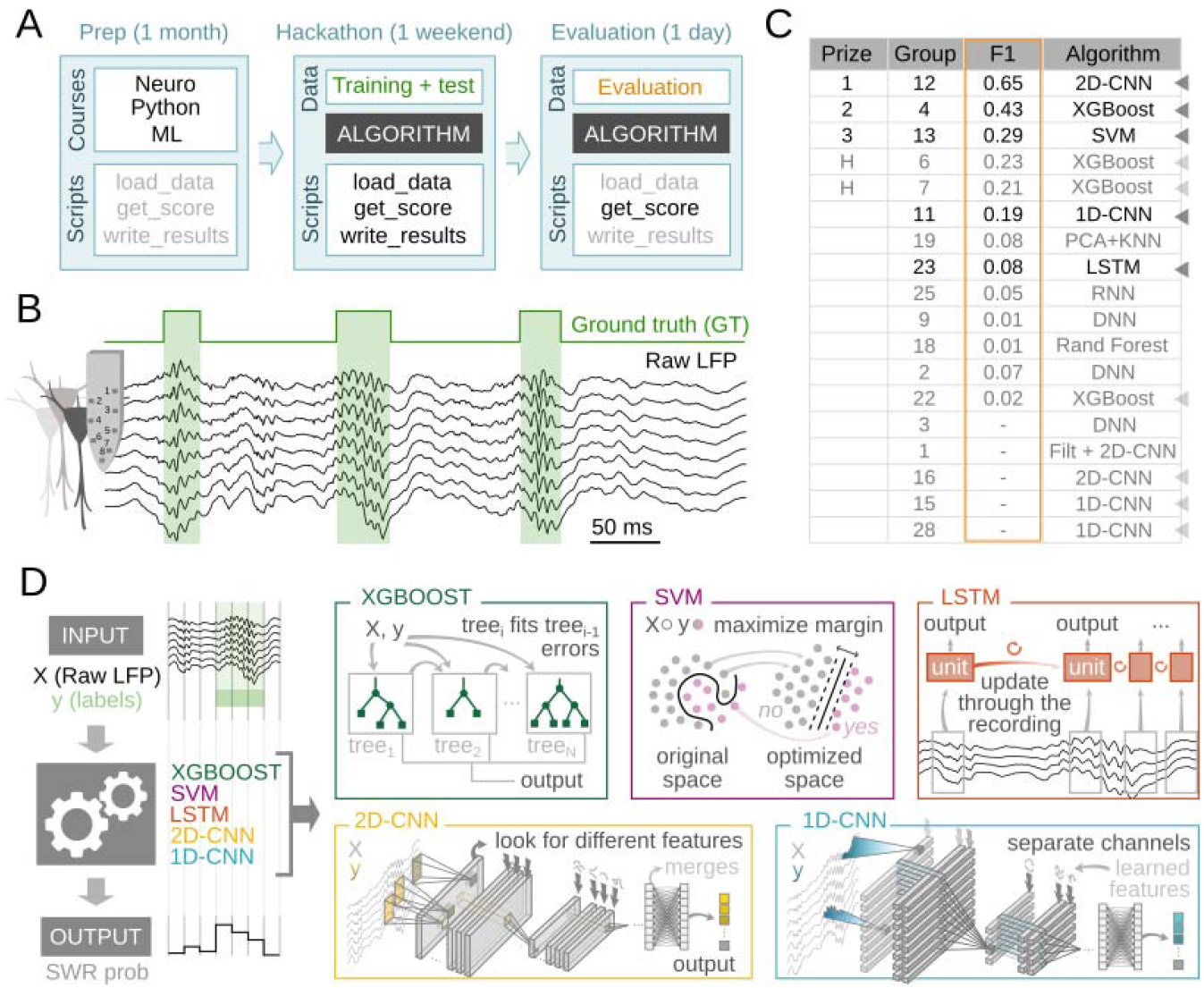
Unbiased community-based proposals of ML models of SWR. **A**, Organization of the hackathon. A preparatory phase (Prep) established the basic grounds of the challenge in terms of minimal knowledge about SWR, Python programming and Machine Learning (ML) models. It also looked to standardize scripts and data management. The second phase consisted on the hackathon, which lasted over 53h during three days, with participants having access to the annotated training dataset and some Python scripts. During the last evaluation phase, a new test set was released to participants 3 hours before the end of the hackathon. Solutions were ranked using the F1-score (see methods). **B**, Example of the training data consisting on 8 channels of raw LFP (black) sampled at 30 kHz, with the manually tagged ground truth (GT), corresponding to SWR events. **C**, Results from the hackathon. Solutions were ranked by the F1-score. F1 represents the harmonic mean between Precision (percentage of good detections) and Recall (percentage of detected GT events). Deep Neural Networks (DNN), Convolutional Neural Networks (CNN), Recurrent Neural Networks (RNN) with/without Long-Short Term Memory (LSTM); Random Forest decision trees (Rand Forest), Extreme Gradient Boosting (XGBoost), Support Vector Machines (SVM), k-Nearest Neighbors (kNN). Chosen solutions are marked with arrowheads. Darker arrows point to the group that got the highest score of each particular architecture; light arrows point repeated architectures. **D**, Schematic representation of the SWR detection strategy and the 5 ML models used in this work.

Participants submitted eighteen different solutions (Fig. 1C). The most used architecture was the Extreme Gradient Boosting (XGBoost; 4 proposals), a decision tree-based algorithm very popular for its balance between flexibility, accuracy and speed^44^ (Fig. 1C). Some other popular architectures were one and two-dimensional Convolutional Neural Networks (1D-CNN, 2D-CNN; 3 and 3 solutions, respectively), Deep Neural Networks (DNN, 3 solutions)^45^, and Recurrent Neural Networks (RNN; 2 solutions)^45^ (Fig. 1C). RNN were presented in both their standard feed-forward version, and as the Long-Short Term Memory (LSTM) version that includes feedback connections, more suited for processing time series data^46^.

Although all these architectures are neural networks typically used for pattern recognition, the way they process and learn from data is remarkably different. For example, whereas CNNs are based on kernels specialized in spotting particular spatially contiguous features of the input, LSTMs use memory cells that look for time-dependent relationships in the data. Two other algorithms were also submitted: a Support Vector Machine (SVM; 1 solution; Fig. 1C) and a clustering-based solution based on dimensionality reduction by Principal Component Analysis (PCA), followed by k-Nearest Neighbors (kNN) clustering (1 solution; Fig. 1C). From the 18 solutions submitted, 5 were not functional and could not be scored (Fig. 1C, bottom). Analysis of the hackathon experience in relationship to the submitted solutions are summarized in Fig. S1 (see methods for details).

**Figure S1.**
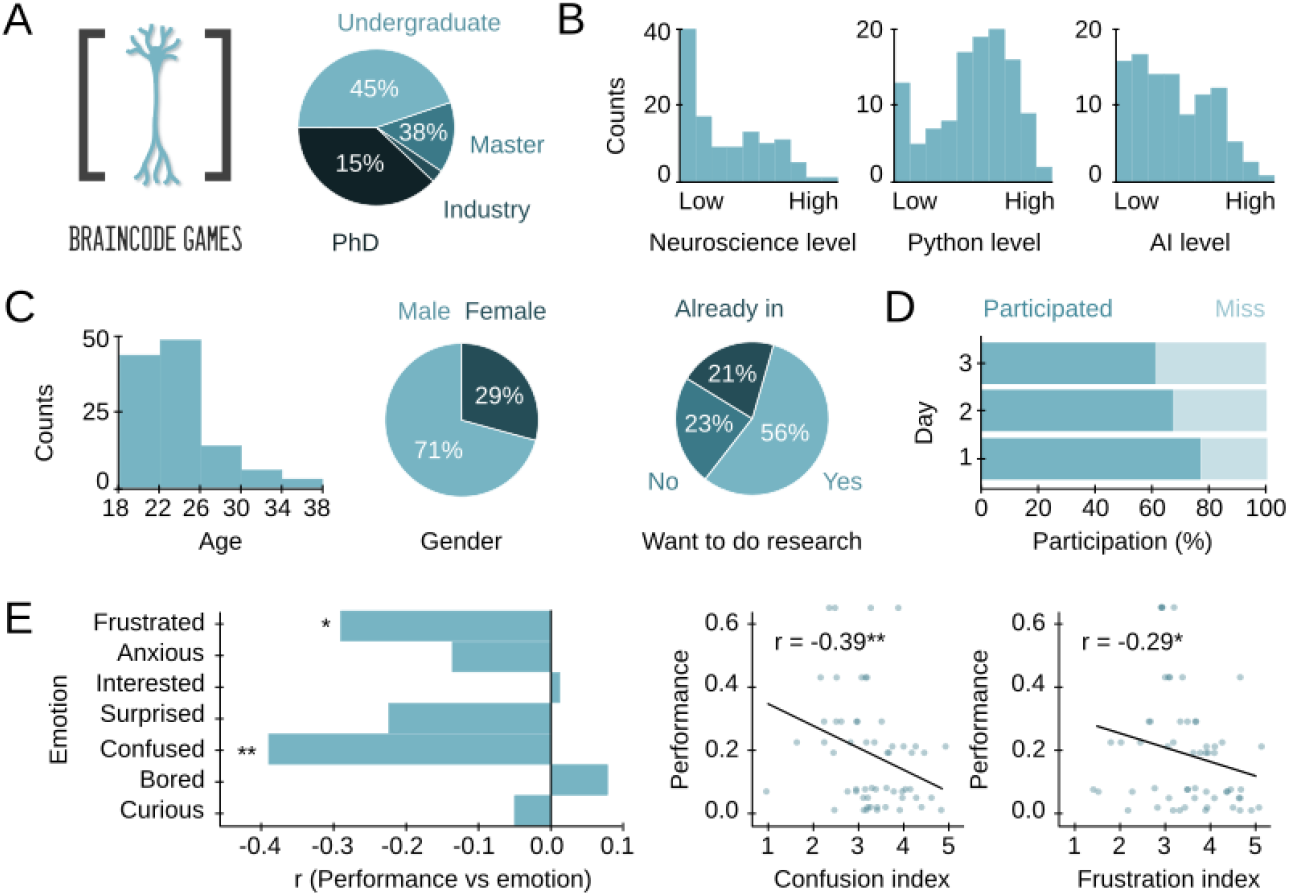
Information about the hackathon. **A**, A hackathon was organized to seek for community-based solutions to the SWR challenge from people unfamiliar to SWR neurophysiology. Among the 116 participants, there were undergraduate students (45%), Master students (38%), PhD students (15%), and industry workers (3%). **B**, There was a general lack of neuroscience knowledge, although most participants declared a high-level performance in Python. Most groups integrated people with programming abilities and basic ML knowledge. **C**, Participant age (left), gender (middle; 71% male, 29% female participants) and involvement in research (right; 21% already in research; 56% interested in doing basic research; 23% not motivated for basic research activities). **D**, Self-reported participation rate during the three days of the hackathon. **E**, Correlation between the performance metric of the proposed solution and emotional states of participants as quantified from their responses to surveys recorded during the hackathon (Spearman rank-order correlation *, p<0.05; **, p<0.01). Only performance of functional solutions were used. See Methods for details.

We sought to identify the more promising architectures for a subsequent in depth analysis. Performance of submitted ML models was measured using the F1-score (see next section). The best performances were achieved by the 2D-CNN, one of the XGBoost models, and the SVM algorithm. Since 1D-CNNs and RNNs were submitted by several groups, and given their previously successful application to SWR detection^28,31^, we decided to include them as well, resulting in five different machine learning architectures (Fig. 1C; dark arrowheads).

The goal of the ML models is to identify the presence of a SWR (or part of it) in a given analysis window (Fig. 1D, left). The selected ML architectures covered a range of processing strategies (Fig. 1D, right). XGBoost is a very popular ML algorithm that uses many decision trees in a parallel fashion, making it one of the fastest algorithms^47^. SVM regression lays within the statistical learning framework, and its objective is to find a new space where samples from different categories (SWRs vs no-SWRs) are maximally separated, making it one of the most robust classification methods^48^. LSTMs are especially suited for regression and classification of temporal series like in natural language processing, using a memory-based strategy to extract relationships between non-continuous time points^46^. CNNs represent a very common approach for many detection and classification tasks applied to different data modalities (1D for signals, 2D for images and 3D for video or volumetric reconstructions), and can approach human performance on many tasks^49^. While 2D-CNNs process input data by considering adjacency on both dimensions (spatial and temporal, in our case), the 1D-CNN solution treats each channel independently and only considers time adjacency, making them two distinct processing algorithms.

This community-based ML architecture bank that was produced by participants who were unfamiliar with SWR studies can be used to evaluate the problem of SWR automatic detection in experimental contexts. We next focused on standardizing processing and retraining the different models.

### Standardization and retraining of selected algorithms

After careful examination of the submitted solutions, we noticed that data pre-processing and training strategies were very different between groups. Data characteristics, like the sampling frequency or the number of channels used for detection can influence operation. To standardize analysis, we chose to down sample to 1250 Hz, and normalize input data using z-scores, which account for differences in mean values and standard deviation across experimental sessions.

We then retrained the submitted ML architectures using the same training set of the hackathon. We randomly divided the dataset into a training set (70%), and a test set (30%) to evaluate their performance in unseen data prior to a more thorough validation (Fig. 2A). We explored a wide range of hyper-parameters for each architecture, which included the number of LFP channels (1, 3 or 8), the size of the analysis window (from 6.4 up to 50 ms) and model-specific parameters like “maximum tree depth” for XGBoost, “bidirectionality” for LSTM or “kernel factor” for CNNs (Fig. 2A). A trained ML architecture set with a particular combination of its hyper-parameters gives rise to a particular “trained model” (Fig. 2A). Because each architecture had different numbers of hyper-parameters, we ended up with different numbers of trained models for each architecture (1944 for XGBoost, 72 for SVM, 2160 for LSTM, 60 for 2D-CNN, and 576 for 1D-CNN). We then used the test set to choose the 50-best models from each architecture, and further tested their performance using a new validation dataset (7586 SWR events; 21 sessions from 8 mice), previously used for the 1D-CNN model^31^ (Fig. 2A, right).

**Figure 2:**
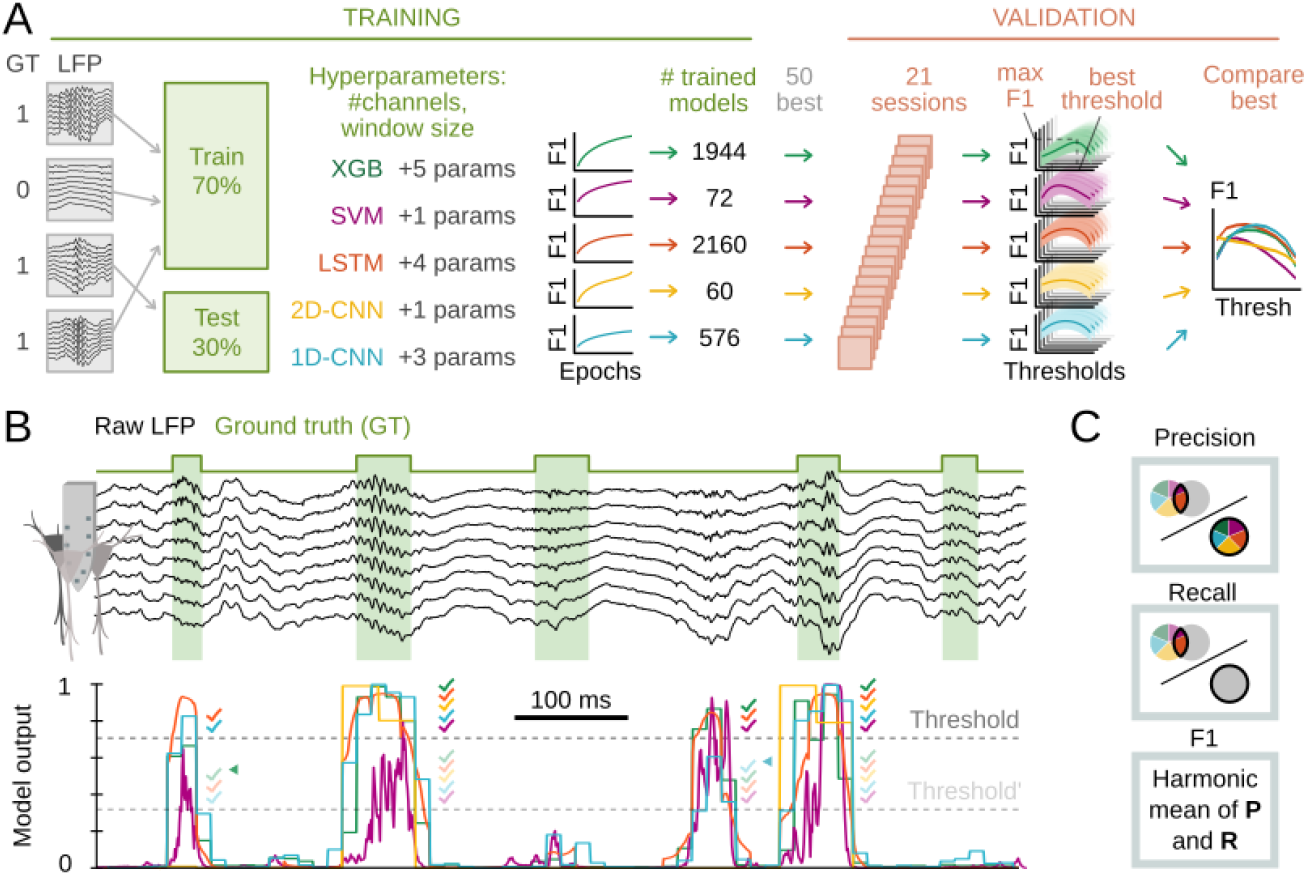
Training design and performance of ML models. **A**, Training and selection criteria scheme. The training dataset used in the hackathon was z-scored and down-sampled to 1250Hz. Training data were shuffled and distributed into train and test subsets (70%-30% respectively). Each architecture was trained to optimize F1 of the test set using several parameters. The 50 best models were tested over a new validation data set (7586 events; 21 sessions from 8 animals), generating an F1 vs threshold curve per model/ architecture. Among these 50, the model with highest mean F1 was selected for between-models comparison (right panel). **B**, LFP example of the validation set and the corresponding model outputs per window of analysis. Note different duration of true events. Setting a threshold allows defining the windows containing detected events. Colored ticks represent detections by the different models. Two different thresholds (dark and light gray) can influence what events are detected. Note how detections marked with arrows are dismissed when the threshold increases. Since SWRs constitute about 1-4% of the total detected events are not computed for performance. **C**, Schematic illustration of Precision (percentage of good detections), Recall (percentage of ground truth events that have been detected) and F1-score (harmonic mean between Precision and Recall).

The goal of training is to make the model output as similar as possible to the ground truth (GT). Because model outputs are continuous numbers between 0 and 1 representing the probability of the presence of the event in the window of analysis, choosing the detection threshold can affect performance (Fig. 2B). Lower thresholds would result in more detections (Fig. 2B, light-gray discontinuous threshold line), normally implying a larger number of both True and False Positives, while higher thresholds are more conservative at the expenses of False Negatives (Fig. 2B, dark-gray threshold line). An ideal model would perform well regardless the threshold, but in practice selecting the threshold that optimizes the True Positive-False Positive trade-off is unavoidable but crucial for experiments. A performance score that takes into account this trade-off is the F1-score, computed as the harmonic mean between Precision (percentage of good detections) and Recall (percentage of detected GT events) (Fig. 2C). F1 values of 1.0 would reflect a perfect match between detections and GT, whereas 0.0 reflects a perfect mismatch. Note this was the same score used to rank models in the hackathon.

After training all architectures by optimizing F1-scores over the test set, we assessed generalization and performance using the validation dataset. We inspected what parametric combinations gave rise to optimal ML models, and found a remarkable variety of distributions (Fig. S2A). All architectures showed a great deal of variability, with almost all available parameter combinations covered. However, some parameters showed biases that depended on the ML architectures, pointing to the necessary requirements for a good performance. For example, all of the 50-best XGBoost models used 8-channels, and in general, more than 1-channel was used across successful architectures (Fig. S2A). Furthermore, different architectures had distinct ranges of parameter values. XGBoost models required longer time windows (25 ms), whereas most SVM models employed shorter windows (<3.2 ms). LSTM, 2D-CNN and 1D-CNNs with variable window sizes all showed very strong performance for >12.8 ms. Finally, LSTM models used both uni- and bi-directional input flow, whereas all of the best models resorted to bidirectionality, suggesting that there should be SWR information also coded in the period preceding an event^50^.

A plug-and-play toolbox to use any of the best 5 models of each architecture for SWR detection is available: https://github.com/PridaLab/rippl-AI.

**Figure S2:**
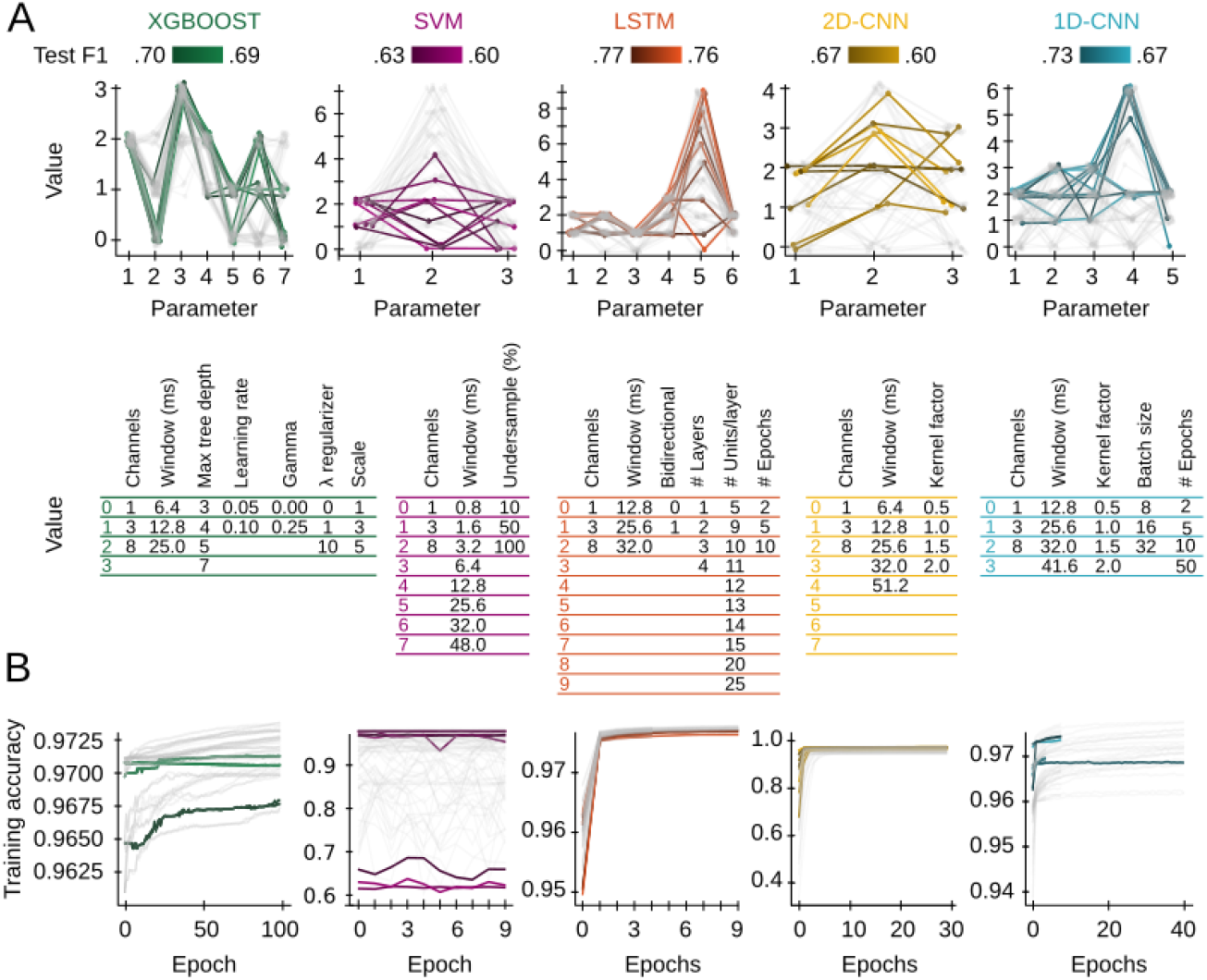
Definition of parameter space in the different ML architectures. **A**, Results from the different architectures in the training dataset: XGBOOST, SVM, LSTM, 2D-CNN and 1D-CNN. Tables indicate the different hyper-parameters used to train each architecture. The resulting 10-best models are color-coded by their F1-score in the validation dataset. The remaining 40-best models are shown in light gray. **B**, Evolution of accuracy along training epochs for the ML models shown in A.

### Influence of the temporal and spatial sampling in training performance

Next, we sought to evaluate the relationship between model performance, parameters and LFP input characteristics. Given the relevance of the temporal and spatial LFP sampling in the definition of SWRs^31^, we started evaluating how the size of the analyzed window and the number of recording channels influenced performance. In order to have as much data as possible, we used F1-scores of all the trained models over the test set.

We found that XGBoost and LSTM were very stable, with performances changing very little for any combination of window size and the number of channels used, suggesting that these architectures can capture SWR features that are relatively invariant across temporal/spatial windows in the input data (Fig. 3A,B). Interestingly, the training parameter that most influenced these two architectures was the number of LFP channels, with 3 and 8 channels providing better performances (Fig. 3A).

**Figure 3:**
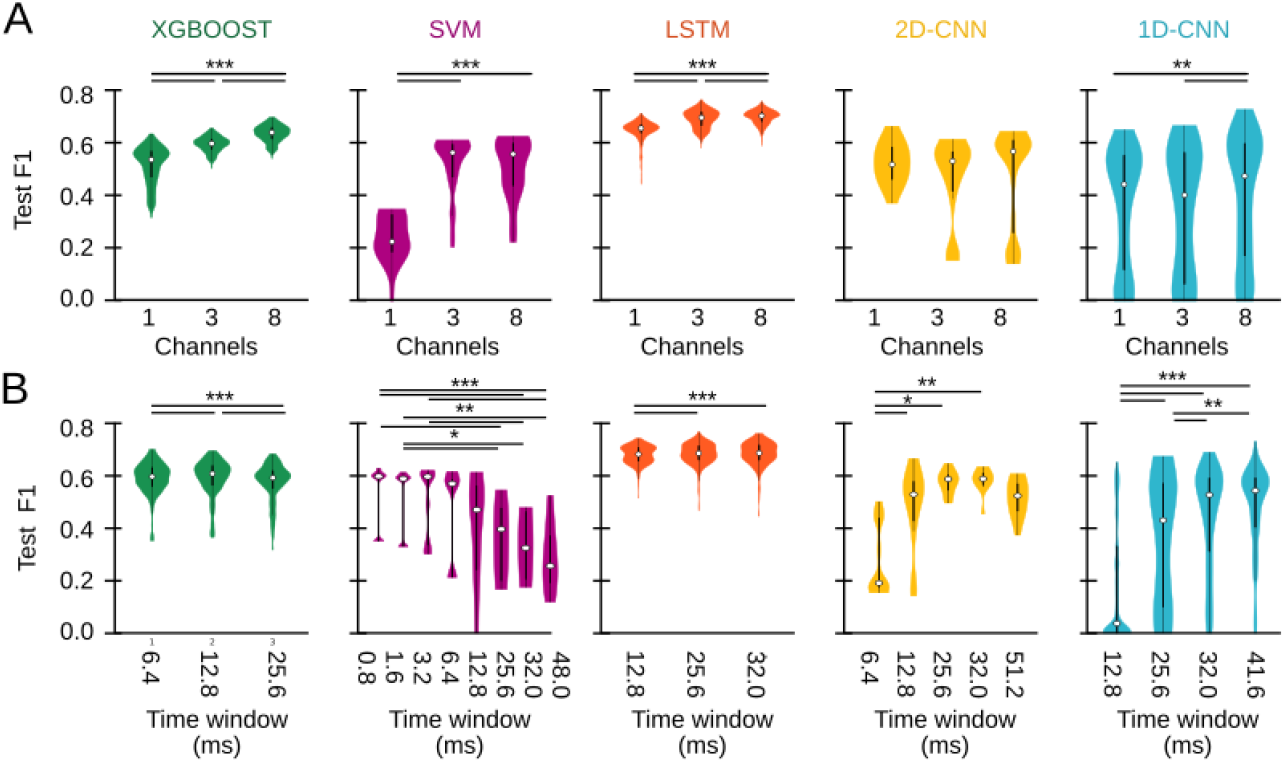
Influence of number of channels and analysis window on training performance. **A**, Final test F1-score of all trained models depending on the number of input channels: one (pyramidal channel; see methods), three (pyramidal channel and extreme channels), or eight (all channels of the probe). Kruskal-Wallis tests with repeated measures for every architecture: XGBOOST, Chi2(2)=1282.2, p<0.0001; SVM, Chi2(2)=33.1, p<0.0001; LSTM, Chi2(2)=964.4, p<0.0001; 2D-CNN, not significant; 1D-CNN, Chi2(2)=14.6, p=0.0007. Post hoc tests *, p<0.05; **, p<0.01, ***, p<0.001. **B**, Same as panel A, but depending on the time window used for analysis. Kruskal-Wallis tests with repeated measures for every architecture: XGBOOST, Chi2(2)=369.5, p<0.0001; SVM, Chi2(7)=48.8, p<0.0001; LSTM, Chi2(5)=48.0, p<0.0001; 2D-CNN, Chi2(4)=16.5, p=0.0024; 1D-CNN, Chi2(3)=126.5, p<0.0001. Post hoc tests *, p<0.05; **, p<0.01, ***, p<0.001.

Spatial information was also important for the SVM model, which scored poorly using a single versus several channels (Fig. 3A; magenta). As mentioned above, temporal resolution was also critical for SVM, which required smaller time windows of <3.2 ms to succeed in detecting SWR (Fig.3B). For analysis windows >6.4 ms (i.e. the temporal scale of one 150Hz-ripple oscillation) performance dropped significantly, indicating that a single SWR cycle and its particular waveform across channels are optimal input information for the SVM architecture to detect events. This effect could be due to the low number of trainable parameters used for SVM (ranging from 1 to 100; see methods), which requires less but more informative data to achieve good performances.

Finally, both the 2D- and 1D-CNN models had similar performance for any number of channels, although there was also a trend for higher spatial sampling (Fig. 3B, yellow and acqua). Interestingly, both CNN models presented a large F1 dispersion because their performance was very dependent on the window size (Fig. 3B). The 2D-CNN model exhibited maximal F1-score for 32ms, while most 1D-CNN models best scored for 25 ms (Fig. 3B). This may be related to the number of training parameters: the more parameters, the more complex tasks these algorithms can solve, provided the amount of training data is representative enough of the expected variance. This supports accurate detection in longer LFP windows. Examination of the remaining parameters suggested additional differences across architectures (Fig. S3A-E). Interestingly, evaluating their impact on F1-scores confirmed the effect of channels and window size on model behavior (Fig. S3F). For CNN models, the batch size (1D-CNN) and the number of kernels (2D-CNN) were also critical.

**Figure S3:**
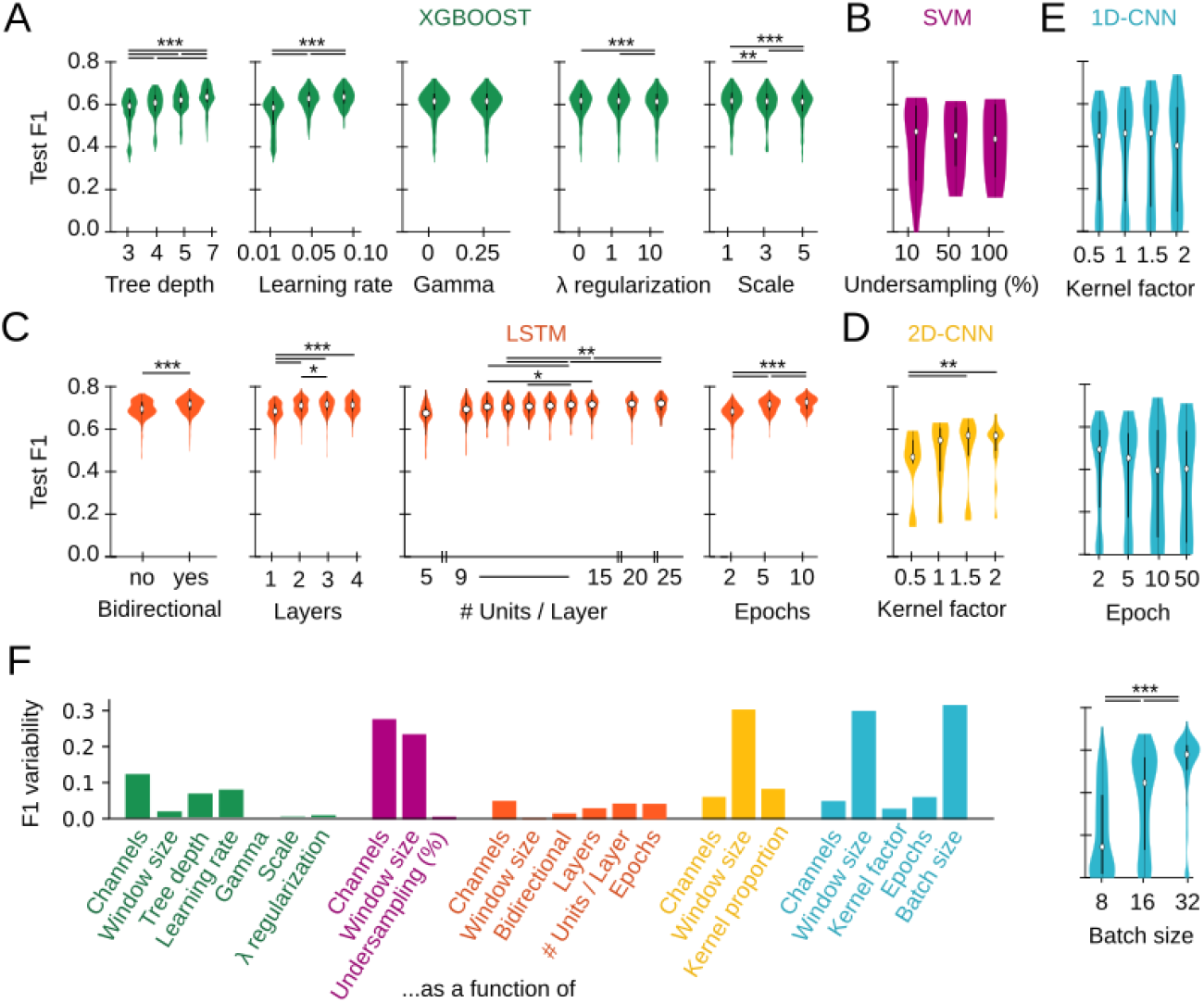
Influence of architecture-specific training parameters on performance. **A-E**, F1-scores from the test set for all models of each architecture. All statistical tests were Kruskal Wallis (KW) with repeated measures. **A**, XGBoost training parameters: maximum tree depth (KW: Chi2(3)=1321.6, p<0.0001), learning rate (KW: Chi2(2)=1109.4.6, p<0.0001), gamma (KW not significant), lambda regularization (KW: Chi2(2)=67.8, p<0.0001) and scale (KW: Chi2(2)=111.6, p<0.0001). Post hoc tests *, p<0.05; **, p<0.01, ***, p<0.001. **B**, SVM training parameters: under-sampling percentage (KW not significant). Higher % of undersampling means training the model with higher representativity of GT data. **C**, LSTM training parameters: bidirectionality (KW: Chi2(1)=320.1, p<0.0001), number of layers KW: Chi2(3)=602.4, p<0.0001), number of units per layer (KW: Chi2(9)=543.8, p<0.0001) and training epochs (KW: Chi2(2)=836.1.6, p<0.0001). **D**, 2D-CNN training parameters: number of kernels scaling factor (KW: Chi2(3)=16.0, p=0.0011), number of epochs and batch size (KW not significant). **E**, 1D-CNN training parameters: number of kernels scaling factor (KW not significant), number of training epochs (KW not significant), and batch size (KW: Chi2(2)=196.9, p<0.0001). **F**, F1-score variability as a function of all training parameters. F1 variability was computed as the difference between the maximum and minimum mean F1.

### Comparison between optimized models

The analysis above provided insights on how input characteristics and processing parameters can influence detection performance in different ML models. Understanding how each architecture learns to identify ripple-like events can not only can aid the development of new tools, but unveil what are the key LFP features used for detection. We thus evaluated conditions for their best performance.

For fair comparison between architectures, we selected the 10-best models from the validation set. Remarkably, our previously published 1D-CNN model^31^ was among the 10-best 1D-CNN, outperforming other configurations. Plotting F1-scores of all models across a range of thresholds allowed visualization of their performance stability as a function of the probability threshold (Fig. 4A).

**Figure 4:**
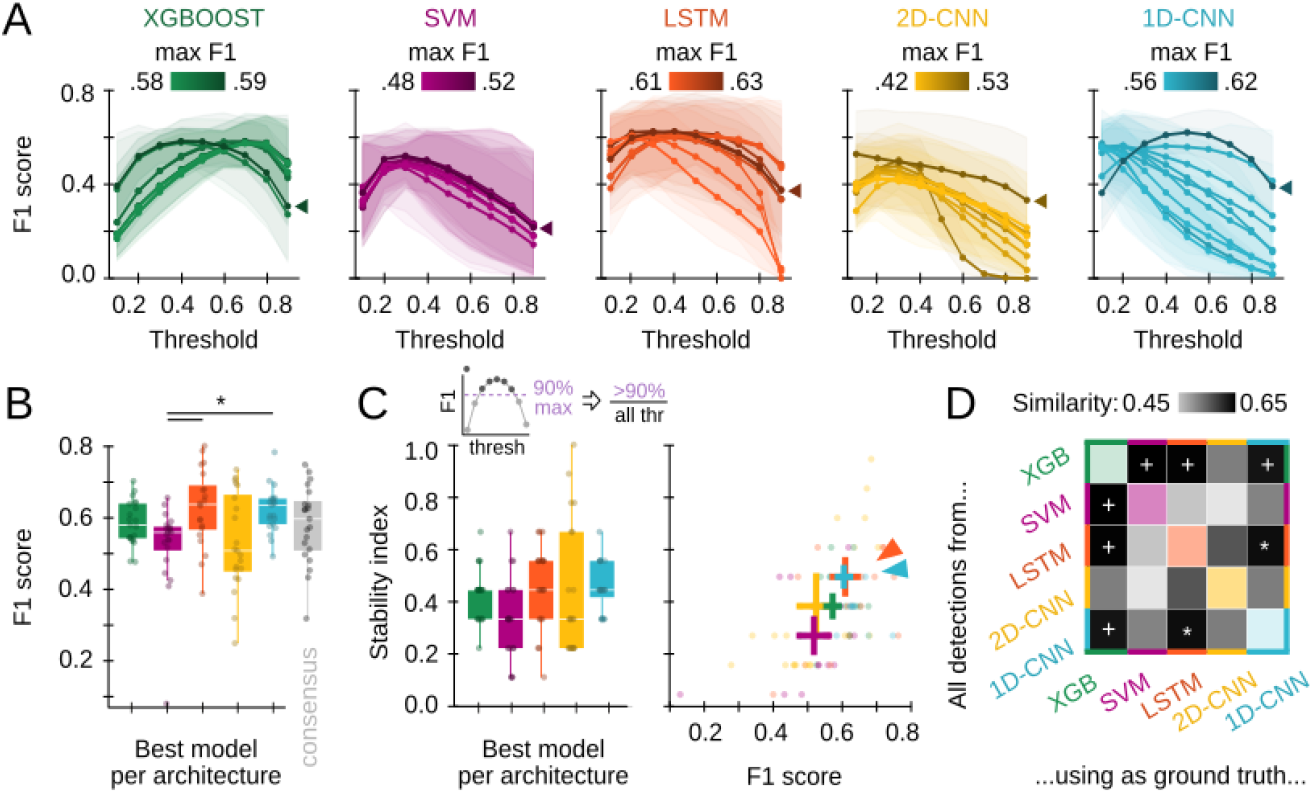
Comparison between best performing ML models. **A**, F1 against threshold from the 10-best models of each architecture as evaluated in the validation set. Each line represents the performance of one trained model, colored by its maximal F1 (mean from all sessions is plotted in dark color). Data reported as mean±95% confidence interval for validation sessions. Arrows indicate the best model of each architecture. **B**, F1-scores for the best model of panel A. Thresholds used are: 0.4 for XGBoost, 0.5 for SVM, 0.4 for LSTM, 0.1 for CNN2D, 0.5 for CNN1D. Each dot represents a session of the validation set (n=21 sessions; 8 mice). In gray, the F1-score for a consensus detector. Kruskal-Wallis, Chi2(5)=26.9, p<0.0001; post hoc tests *, p<0.05; **, p<0.01, ***, p<0.001. **C**, Stability index for the best model of each architecture (left), and the stability index vs the F1 (right). Kruskal-Wallis, Chi2(4)=10.5, p=0.03; post hoc tests. **D**, Similarity between predicted events of different architectures. Models are the same as in panels B-C. To measure the similarity, the mean F1 across validation sessions have been computed, using detected events in the y-axis as detections, and detected events in the x-axis as ground truth. Note the similarity between LSTM and 1D-CNN (white *), and that by XGBoost against SVM, LSTM and 1D-CNN (white +).

We analyzed their performance along a range of characteristics (performance, robustness, and threshold dependency) to better inform their selection depending on research applications. Five of the 10-best trained models of all architectures are available at https://github.com/PridaLab/rippl-AI/blob/main/optimized_models/

The consistency of F1-threshold curves depended on the model architectures (Fig. 4A). Most models reached their maximal F1-score at relatively low threshold values of 0.3-0.4 and remained stable until a probability of around 0.5-0.7. Such a behavior indicates robust performance, since even low probability (i.e., relatively uncertain) output predictions overlapped with the ground truth. This property is very useful for online experimental applications, when choosing different thresholds is not manageable, making detection more robust. Interestingly, we found that XGBoost models exhibited good performance at two threshold ranges (0.2-0.4 and 0.6-0.8), depending on how trained models penalized False negative predictions. Similarly, for both CNN architectures, we found several models operating sharply at low thresholds, while others exhibited a relatively stable operation in the 0.4-0.6 range especially for 1D-CNN models. We confirmed the variability of different models within a given architecture by looking at their Precision vs Recall curves for the entire threshold range (Fig. S4A). This variability suggests that even when arising from the same architecture, algorithmic processes and detection strategies by which the different models were detecting SWR events could differ. This may provide a range of models for different applications.

Next, we selected the model that reached the highest F1 value from each architecture (Fig. 4A, best models, arrowheads), and compared their scores using all validation sessions (Fig. 4B). We found that the LSTM and 1D-CNN best models outperformed other architectures, with mean F1-scores over 0.6 (as a reference, the inter-expert F1-score in our lab is ∼0.7^31^. Precision-Recall curves from these two models clearly stood out of the other solutions (Fig. S4B). Importantly, a consensus prediction based on the 5-best models did not perform better than individual architectures alone (Fig. 4B; gray).

Given the importance of consistent threshold performance for practical applications, we quantified the robustness of F1-threshold curves for the best models using a stability index in the validation dataset (see methods). Models with a stability index of 1.0 provide at least 90% of its maximal performance for any threshold value, a property especially suitable for experimental applications. While the best 2D-CNN model exhibited stability in some validation sessions, the best LSTM and especially the best 1D-CNN best models exhibited more consistent behavior (Fig. 4C, left; Fig. S4C). We confirmed this result by plotting the stability index versus F1, where both the best LSTM and 1D-CNN best models clearly segregated (Fig. 4C, right; arrowhead).

**Figure S4:**
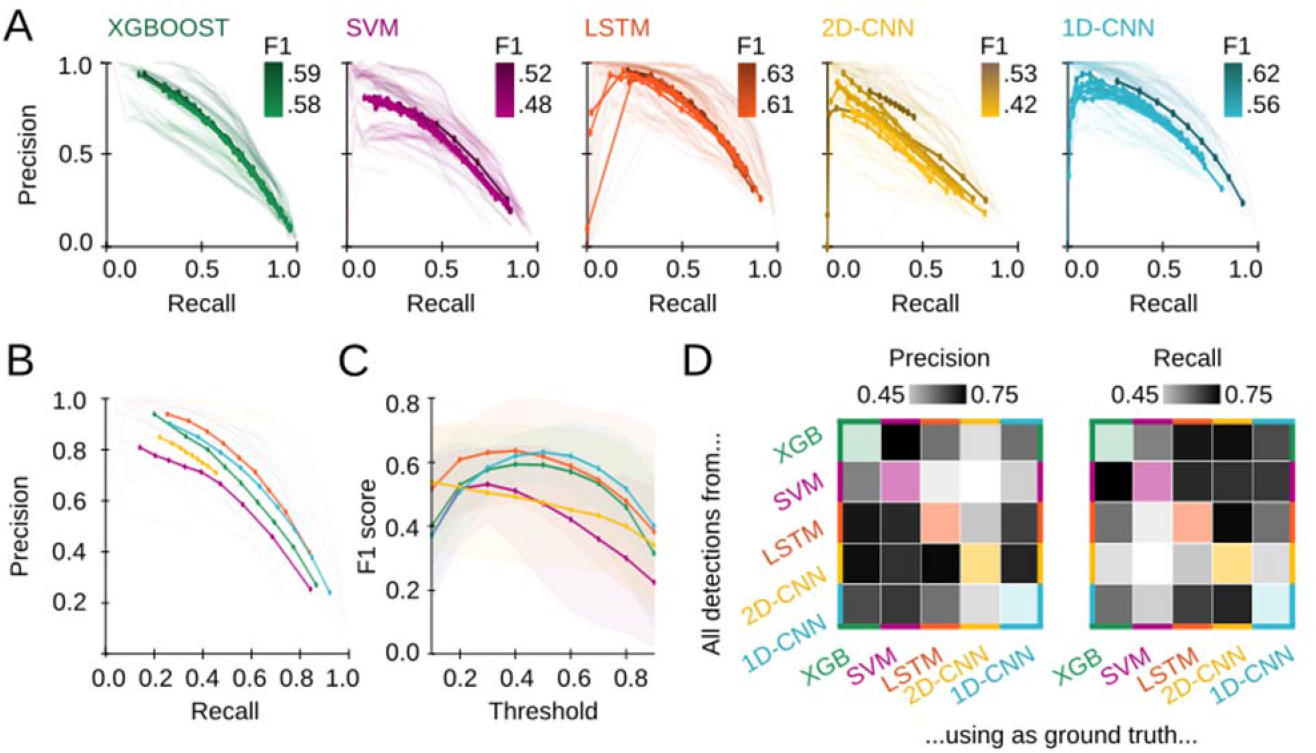
Precision-Recall curves of optimized models. **A**, Precision (P) vs Recall (R) curves for the 10-best models of each architecture. Each dot represents P-R values for a particular threshold. Each line represents the performance of one trained model, colored by its maximal F1 (mean of all sessions is plotted in dark color; sessions are light colored). **B**, P-R curves for the best model of each architecture (all thresholds). Thick lines represent mean values. Thin lines curves are individual validation sessions. **C**, F1-score as a function of the threshold. Data reported as mean±95% confidence interval for validation sessions. **D**, Similarity between the events predicted by the best model (maximum F1) of each architecture. Models shown are the ones with maximum F1. To measure the similarity, we computed the mean Precision (right) and Recall (left) across validation sessions have been computed, and used detected SWR events of models in the y-axis as detections, and detected events of models in the x-axis as ground truth.

Finally, to evaluate whether the different models were targeting similar or different subsets of SWR events, we compared how similar their detections were. To quantify this similarity, we computed the F1 between both groups of detections, using one of them as the ground truth (Fig. 4D). Interestingly, the 1D-CNN and LSTM showed a high level of similarity, in line with their consistent and accurate behavior (Fig. 4D, white *). XGBoost scored a high similarity with all other architectures except for the 2D-CNN (Fig. 4D, white +). Possibly, this reflects the fact that very few of the XGBoost detections were also predicted also by 2D-CNN, leading to a very low Precision (Fig. S4D). In general, high similarities did not seem to be caused by a particularly high Precision or Recall (model A detects so few events that all coincide with detections of model B), but by a good balance between both (events of model A and B highly overlap) (Fig. S4D).

### Effect of different ML models on the features of detected SWRs

Results above suggest that different models may be relying on different strategies for recognizing SWRs. We thus wondered whether models could be biased towards SWRs with different features (e.g. frequency, amplitude, etc…), and whether these biases could also be reflected over different ranges of output probabilities.

In order to evaluate these issues, we resorted to a low-dimensional analysis of SWRs which allows for their unbiased topological characterization^14^. In this strategy, SWR events are considered points in an N-dimensional space, where each dimension X (dimX) represents the LFP value sampled at a given timestamp X (Fig. 5A). In our case, as events were GT ripples of 50 ms sampled at 1250Hz (i.e. 63 timestamps), the original space was 63 dimensions. Plotting all SWR events will result in a point cloud, with events sharing similar LFP features lying close to each other, while those of different characteristics distribute separately (Fig. 5A). To ease visualization, the SWRs were embedded in a low-dimensional representation using Uniform Manifold Approximation and Projection (UMAP)^14,51^.

**Figure 5.**
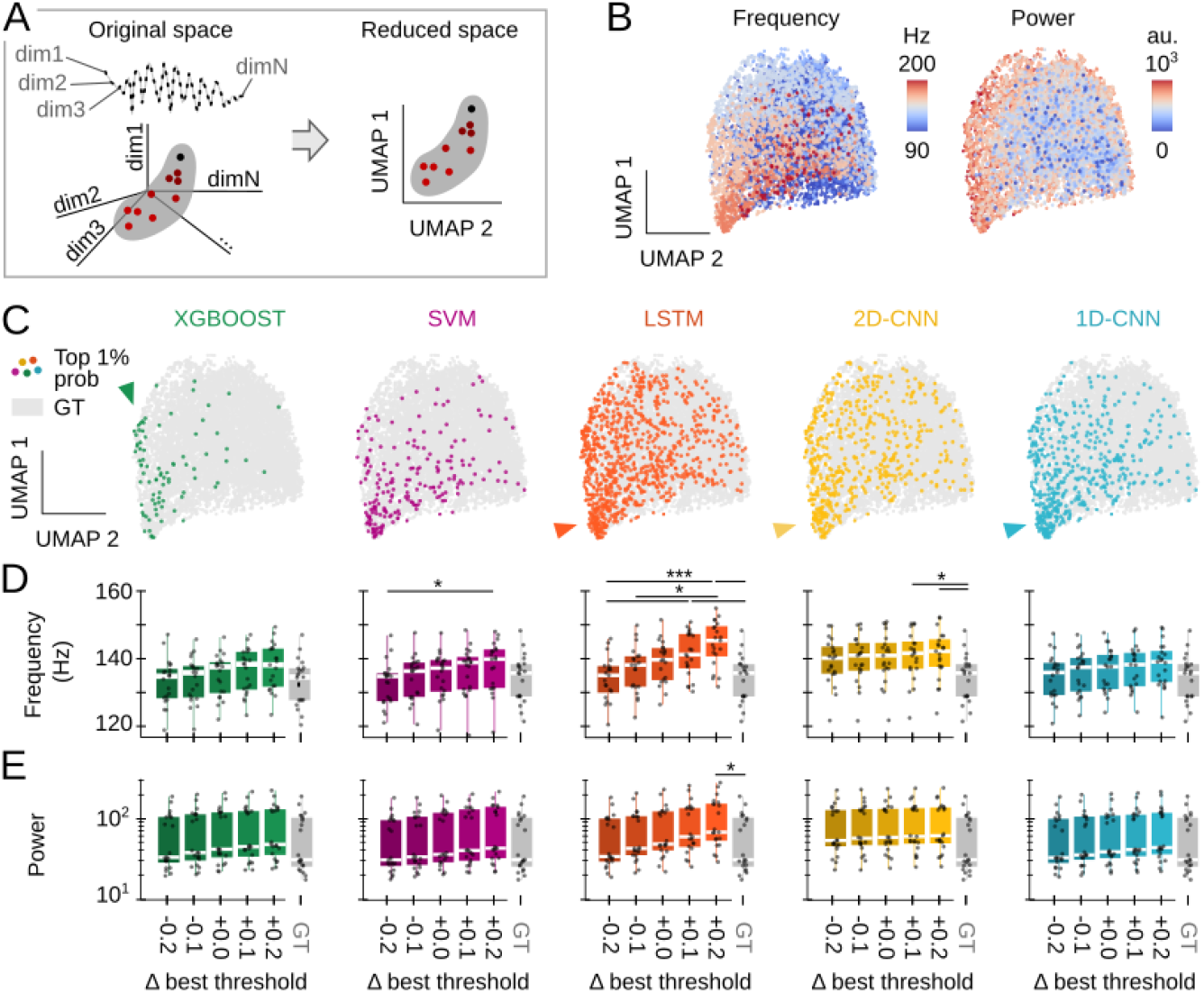
Effect of ML models and thresholds on the type of detected SWR. **A**, Low-dimensional analysis of SWR features^14^. GT ripples are represented into a high-dimensional space by mapping each timestamp to a particular dimension. Since the sampling rate is 1250Hz, and windows around SWRs were cut to 50ms, there are 63 timestamps per event, and so the original space has 63 dimensions. The SWR cloud is embedded in a low-dimensional space using UMAP. **B**, UMAP embedding projected into the two first axes. Each dot represents a GT ripple, and its color reflects its frequency (left) and power (right). Note how ripples in the cloud are distributed according to frequency and power, meaning that in the original space ripples with similar features are close together. **C**, Colored dots superimposed over gray GT data represent the top 1% of detected events for every given architecture, i.e., True Positive events with an output SWR probability above 99% of the maximum probability for that given model. Note that different distributions of events in the cloud reflect biases of ML model used for detection. **D**, Frequency of True Positive SWR detected by each architecture. Each dot represents the mean frequency of detected ripples of one validation session (21 sessions from 8 animals). Kruskal-Wallis tests for every architecture: XGBOOST, not significant; SVM, Chi2(5)=11.1, p=0.049; LSTM, Chi2(5)=29.9, p<0.0001; 2D-CNN, Chi2(5)=13.8, p=0.017; 1D-CNN, not significant. **E**, Spectral power of True Positive events detected by each architecture. Kruskal-Wallis tests for every architecture: XGBOOST, SVM, 2D-CNN and 1D-CNN are not significant; LSTM, Chi2(5)=14.0, p=0.016. Post hoc tests *, p<0.05; **, p<0.01, ***, p<0.001.

First, we analyzed how ripple frequency and power were distributed in the UMAP embedding by coloring each dot (i.e. each SWR) based on their frequency (Fig. 5B, left) and power (Fig. 5B, right). As expected from our previous work^14^, these features followed different distributions, segregating high-frequencies towards the bottom of the cloud and high-power events radially out (Fig. 5B). We then inspected events detected by the best model of each architecture by plotting the top 1% detections, defined as True Positive events for which the model output probability was >99% of its maximum probability (Fig. 5C). Interestingly, each model showed different distributions of preferred SWRs. For example, XGBoost was biased towards a subset of high-power and fast SWR events (Fig. 5C, green arrowhead), whereas the SVM model exhibited a more heterogenous distrinution. In turn, LSTM and both CNNs assigned higher probabilities to events that had a good frequency-power balance (Fig. 5C, orange, yellow and blue arrowheads). Note how these models have more colored events, consistent with their higher stability indices reported above (Fig. 4C).

To quantify detection biases in each ML model, we analyzed the frequency and power of their True Positive events and compared them against those in the GT. Consistent with the UMAP distributions, SWR frequency was highly dependent on the threshold for SVM, LSTM and 2D-CNN algorithms (Fig. 5D). The case of LSTM was particularly striking with differences accumulating for all thresholds. Instead, for the SVM and 2D-CNN biases were significant only when thresholds differed ±0.2 from the optimal value (Fig. 5D). As previously reported^31^, the 1D-CNN exhibited roughly consistent behavior with SWR features not statistically different from GT events. SWR power exhibited no major dependency on the threshold in any of the models but the LSTM, especially at higher detection thresholds (Fig. 5E).

Altogether, this analysis suggests that the different ML models can be exploited to detect a wide range of SWRs with different characteristics.

### Using the toolbox to identifying SWRs in non-human primates

A major motivation of our study is to develop methods which can be generalizable for a wider range of detection contexts, including a greater range of species and biomedical applications. Thus, we applied our ML models to LFP recordings from the hippocampus of the macaque, which shares a high level of genetic, morphological and physiological characteristics with that of its fellow primate, the human, while enabling precise localization of signals roughly comparable to those used for the algorithm development. To accomplish this, we recorded hippocampal LFP signals from a freely moving macaque using a multichannel linear probe^52^ (Fig. 6A). Unlike the original high-density probes (20 μm), recordings were obtained every 90/60 μm and spanned CA1 layers (Fig. 6A). As in mice, SWRs were manually identified (4133 events) to generate the annotated ground truth (Fig. 6B). Consistent with the literature^16,17^, macaque SWRs had lower frequencies and higher power as compared to mouse ripples (Fig. 6C).

**Figure 6.**
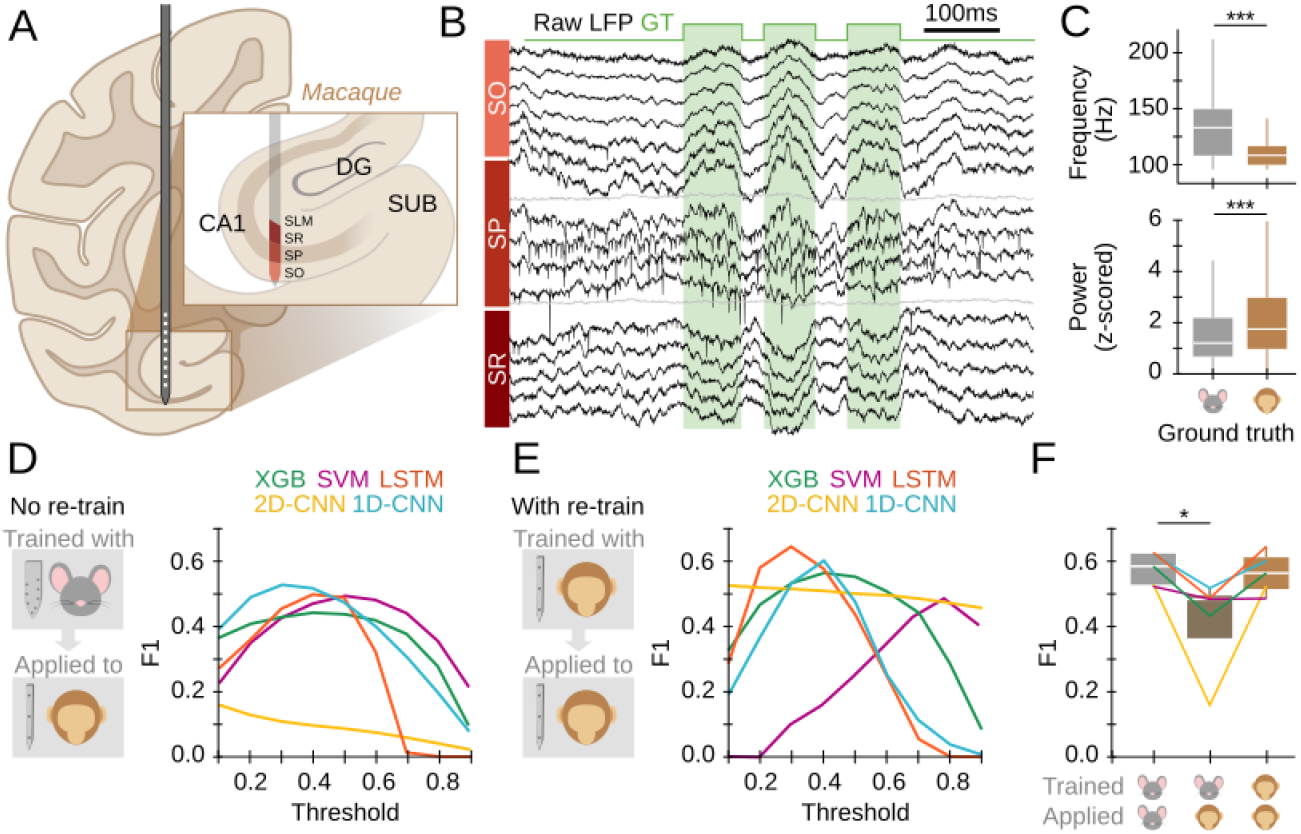
Extending sharp-wave ripple detection to non-human primates. **A**, Linear multichannel probes were used to obtain LFP recordings from the anterior hippocampus of a freely moving monkey. **B**, SWR events were manually tagged (4133 events) as in mouse data. **C**, Significant differences between SWR recorded in mice and monkey. Kruskal-Wallis Chi2=1649, p<0.0001 for frequency; Kruskal-Wallis Chi2=407, p<0.0001 for power. Posthoc, ***, p<0.001. Data from the GT in both cases. **D**. The best model of each architecture trained in mouse data was applied to detect SWRs on the macaque data. Input data consisted of 5 LFP channels of SO, SP and SR, and 3 interpolated channels (see methods for details). We evaluated all models by computing F1-score against the ground truth (GT). Note relatively good results from non-retrained ML models. **E**, Results of model re-training using macaque data. Data were split into a training and test dataset (50% and 20% respectively), used to train the models; and a validation set (30%), used to compute the F1 (left panel). **F**, F1-scores for the maximal performance of each model before and after retraining. Kruskal-Wallis test, Chi2(2)=8.06, p=0.018. Post hoc tests *, p<0.05.

We applied the best model of each architecture trained in head-fixed mice to macaque recordings, and evaluated their performance. For fair comparison, we flipped laminar LFP signals upside down and sampled the channel combination that best matched the characteristic mouse LFP profile (see Methods and layer orientation in Fig. 6A). Strikingly, 4/5 models reached a maximum F1 of ∼0.5 (Fig. 6D), close to their maximal performance on mice data (∼0.6). SVM, 1D-CNN and LSTM exhibited the best performance, as compared to XGBoost and 2D-CNN (Fig. 6D). Importantly, the fact that both LSTM and 1D-CNN have relatively good generalization capability, suggests that they successfully capture shared features of SWRs from mice and macaques.

We next chose to re-train the 5 models with the macaque dataset, using 50% for training and 20% for testing. The remaining 30% was used for validation to compute the final F1. For re-training, we reset all trainable parameters (internal weights) but kept all architectural hyper-parameters fixed (number of input number of channels, input window length, number of layers, etc…). Performances improved after retraining for 4/5 models, reaching a F1 increase of +0.3 for 2D-CNN (Fig. 6E). The best model was LSTM, followed by 1D-CNN and XGBoost. SVM was the only model that did not improve after retraining, but exhibited a shift towards larger thresholds. Furthermore, performance of macaque SWR detection after re-training reached the mouse level (Fig. 6E), suggesting that these models identified similar key features in both species, and could readily be trained to similar levels of accuracy across mice and monkeys. A user-friendly open python notebook to re-train any of the 5 models and use it for event detection is available at https://github.com/PridaLab/rippl-AI/blob/main/examples_retraining.ipynb

## Discussion

Here, we provide a pool of models for automatic SWR detection based on different ML architectures. These include some of the most used ML solutions, such as XGBoost, SVM, 1D- and 2D-CNN and LSTM. The models, which resulted from unbiased community-based proposals, are able to capture a wealth of SWR features recorded in the dorsal hippocampus of head-fixed mice. When applied to LFP recordings from a freely moving macaque, these models were able to generalize detection.

The need for detecting and classifying high-frequency oscillations such as SWR has accelerated over recent years for advanced biomedical applications^28,33,35,41,53^. Identification of these events can help to delineate normal from pathological epileptogenic territories^18,54,55^, and to develop closed-loop intervention strategies for boosting memory function^33,35^. However, spectral-based methods have revealed suboptimal and the community is actively seeking for novel feature-based strategies. Recently, solutions based in ML methods have started to emerge^25,28,31,54^. Using these tools will drive advances not only in online detection of SWRs, but also their unbiased categorization for better mechanistic understanding^11,13,31,56,57^, including their functional ties to visuospatial and episodic memory^10,11,16,34,38,39^.

Amongst the 5 ML models examined here we found the LSTM and 1D-CNN to provide the best performance and reliability using rodent data. The other models exhibited roughly similar behavior depending on the input parameter selection (recording channels and analysis windows). While in general, we found that all of them performed better with high-density multi-channel recordings (8 channels), some of them (e.g. 2D-CNN) exhibited similar results while operating over data sampled with 1 to 3 channels. This suggests they may be able to identify characteristic features with reduced spatial information, which could facilitate applications to human recordings^19,37^.

Detection of SWR candidates with ML models is based on using a probability threshold. We found that the different models exhibited a degree of sensitivity to threshold selection, with LSTM, XGBoost and 1D-CNN providing a wider range of operational stability. This suggests there is a larger range of thresholds in these models which provide relatively similar performance. Instead, SVM and the 2-CNN better operate in a very narrow threshold range. This is very important for online applications, when threshold selection can affect experimental results in real time^25^.

The different ML models are biased towards SWRs with slightly different properties, probably reflecting their internal representations of SWR characteristic LFP features^31^. During training, each model learns to identify what specific LFP features made ripples distinguishable from background LFP signals, so that during SWR detection, the presence of those features raises their output probability. The fact that the properties of detected SWR depend on the probability threshold for SVM and 2D-CNN suggests that frequency and relative power are some of the LFP features these models identified during training. On the contrary, XGBoost, LSTM and 1D-CNN models, which showed less bias, may be capturing other LFP features such as the spatial profile. This is consistent with results from the analysis of the influence of spatial sampling in training performance in these ML models.

When applied to data from the macaque anterior hippocampus, we found that models trained with LFP signals from the dorsal hippocampus of mice can perform relatively well, especially considering established differences in frequency and in LFP shape in monkey and human^10,16,17^. After re-training, their operation improved significantly, reaching the inter-experts’ performance levels at 0.7^31^. This demonstrates the strong capability of the ML models to generalize and suggests the existence of shared features across species. This is of particular importance, because many human applications may not have the exact spatial localization or the same electrode types, in some cases even within studies, and so any effective ML applications will need a high degree of generalizability. It also demonstrates the proof of principle for applying to a wider range of measurements, including other animal models and ripple-adjacent pathologies such as MTL seizures^54^.

More testing along these lines will identify the extent of generalizability across different permutations of species, location, electrode sampling and type, to find the limits of these ML models. To enable such developments, we made several of the 10-best trained models and our coding strategies for detection and retraining openly available to the research community at https://github.com/PridaLab/rippl-AI. They can be tested through open-source notebooks that are ready to use, with enough examples to illustrate their operation capability. Although the notebooks provide easily readable code, they may not be optimal for further code development. That is why the core functions are written as separate Python modules. Users can test these models for SWR detection by loading their own data and defining the channels. The ripple_AI repository has a wide variety of SWR detection tools that include optional supervised detection curation, and a graphical user interface for a quick visual exploration of detected events depending on the threshold chosen, as well as the option of retraining a model with the user’s own data.

This collection of resources joins to the many other community-based approaches for model benchmarking^30,41,o53,58^. Crowdsourced solutions are becoming a tool to advance solutions of particularly difficult problems which require knowledge integration^40,43^. This provides the field with a set of platforms for detecting events from diverse datasets using traditional and state-of-the-art algorithms (e.g., our own ripple-AI toolbox, and https://www.sharpwaveripples.org/). Our toolbox goes beyond SWR detection, easing development of personalized ML models to detect other electrophysiological events of interest^32^. This may be critical in experimental and/or clinical cases, where other detection criteria, i.e. F-values, than those maximizing performance may be more important. For instance, different experiments may call for avoiding either type I or type II errors, and hence the balance between Precision and Recall. Such a versatility of our toolbox may be further exploited to accelerate our understanding of hippocampal function and to support the development of biomedical applications.

## Methods

### The hackathon

In order to explore a wide variety of ML solutions to the problem of SWR detection, we organized a hackathon (https://thebraincodegames.github.io/index_en.html). We specifically targeted people unfamiliar with SWR studies, who could provide unbiased solutions to the challenge. A secondary goal of the hackathon was to promote their interest and engagement at the interface between Neuroscience and Artificial Intelligence especially for future young scientists. The event was held in Madrid in October 2021, using remote web-platforms. Some of us (ANO) coordinated the event. Consent to participate and to share relevant personal data was obtained prior to the event. All participants were informed on the goal of the hackathon and agreed that their solutions were subject to subsequent investigation and modification.

The hackathon comprised 36 teams of 2 to 5 people (71% males, 29% female), for 116 participants in total. They represent 45% of Undergraduate students, 38% Master students, 15% PhD students and 3% non-academic workers (Fig. S1A). On average, they were young in their professional career with 77% of participants being research-oriented (Fig. S1A). Previous to the hackathon, we monitored the participants’ self-declared knowledge level on Neuroscience, Python programming and ML in general using a survey (Fig. S1B). To provide a homogenous floor to address the challenge, we organized three online seminars to cover each of the three topics one month before the activity. Seminars were recorded and made available for review along the experience.

The hackathon was held during one weekend (Friday to Sunday), during which groups had to design and train a ML algorithm to detect SWRs. To standardize the different algorithms for future comparison, they were given Python functions to load the data, compute a performance score, and write results in a common format. Data sets were available from a public research-oriented repository at Figshare (https://figshare.com/authors/Liset_M_de_la_Prida/402282). Participants were given a training set to train their algorithms, and a test set to run validation tests. Data consisted on raw 8-channel LFP signals from the hippocampal CA1 region, recorded with high-density probes, which was used before for similar purposes (Navas-Olive et al. 2022). SWR were manually tagged to be used as ground truth (training set: 1794 events, two sessions from two mice; test set: 1275 events; two sessions from two animals). Since participants had two days to design and train solutions, groups were allowed to interact with us to ask for technical questions and clarification.

We monitored participant’s engagement throughout the hackathon using short questionnaires. This allowed us to check their motivation and other emotional states (i.e., frustration, interest, etc…). Some people dropped out along the days of the hackathon (Fig. S1D). We found many participants felt confused and frustrated with the challenge, and this correlated with their performance, as a posterior analysis suggested (Fig. S1E).

### Datasets and ground truth

Participants of the hackathon were provided with an annotated dataset consisting of raw LFP signals (8-channels) sampled at 30,000 Hz. SWR events were manually tagged by an expert who for each event identified their start and end. The start of the SWR was defined near the first ripple of the sharp-wave onset. The end of the event was defined at the latest ripple or when sharp-wave resumed. The training set consisted of two recording sessions from 2 mice (Navas-Olive et al., 2022). They contained 1794 manually tagged SWRs. The test set consisted of two recording sessions from another 2 mice and contained 1275 SWR events.

For posterior analysis of the results of the hackathon, we used a validation dataset consisting on the 2 test sessions mentioned before plus another 19 sessions for a total of 21 sessions from 8 different mice. They all contained a total of 7423 manually tagged SWR.

The ground truth, i.e. the analysis windows containing a SWR event, was generated for all sessions with the help of a Matlab 2019b tool that considered the window size.

### ML models specifications

Five architectures were selected out of the 18 solutions submitted to the hackathon: XGBoost, SVM, LSTM, 2D-CNN and 1D-CNN. For the purpose of fair comparisons, they were retrained and tested using homogenized pre-processing steps and data management strategies (see below).

We used Python 3.9.13 with libraries Numpy 1.19.5, Pickle 4.0 and H5Py 3.1.0. To build the different neural networks, we used the Tensorflow 2.5.3 library, with Keras 2.5.0 as the application-programming interface. XGBoost 1.6.1 was used to train and test the boosted decision trees classifiers. Scikit-learn 1.1.2 and Imbalanced-learn 0.9.1 were used to train support vector machine classifiers. Analysis and training of the models were conducted on a personal computer (i7-11800H Intel processor with 16 GB RAM and Windows 10).

### Data preparation

For subsequent training and analysis of the architectures selected from the hackathon, all data was pre-processed. From each recording session two matrices were extracted, X, with the raw LFP data, shaped (# of timestamps, # of channels) and Y, the ground truth generated from the expert tagging (# of timestamps). A timestamp of Y is 1 if a SWR event is present.

Values for matrix X were subsampled at 1250 Hz, taking into consideration that SWRs are events that have frequencies in the range of 150 to 250 Hz. Before retraining the algorithms, data was z-scored with the mean and standard deviation of the whole session.

### Training, validation, and test split

For retraining the architectures, the same training dataset provided in the hackathon was used (2 sessions from 2 mice; 1794 SWR events). For initial testing, these two sessions were split according to a70/30 train/validation design. To evaluate the generalization capabilities of the models when presented with unseen data, we used several validation sessions, which provide the necessary animal-to-animal, as well as within animal (sessions) variability. Validation sessions included the 2 test dataset provided in the hackathon and 19 additional sessions (21 sessions from 8 mice, 7423 SWR events)

For retraining, the two training sessions were concatenated and divided into 60 seconds epochs. Each epoch was assigned randomly to the train or validation set, following the desired split proportion. The data was reshaped to be compatible with the required input dimensionality of each architecture (see below). In order to evaluate model performance, two different datasets were used: the test set described above (used for an initial screening of the 50-best models for each architecture), and the validation set (used for generalization purposes).

Identification of SWR events in the data was implemented using analysis windows of different sizes. To identify SWR events detected by the ML models, we set a probability threshold to identify windows with positive and negative predictions. GT was annotated in the different analysis windows of each session. Accordingly, predictions were classified in four categories: True Positive (TP), when the prediction was positive and the GT window did contain a SWR event; False Positive (FP), when the prediction was positive in a window that did not contain any SWR; False Negative (FN), when the prediction was negative in a window with a SWR; and True Negative (TN) when the prediction was negative and the window did not contain any SWR event.

If a positive prediction had a match with any window containing a SWR it was considered a TP, or it was classified as FP otherwise. All true events that did not have any matching positive prediction were considered FN. Negative predictions with no matching true events windows were TN.

With predicted and true events classified into those four categories, there are three measures than can be used to evaluate the performance of the model. Precision (P), which was computed as the total number of TPs divided by TPs and FPs, represents the percentage of predictions that were correct.

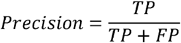

Recall (R), which was calculated as TPs divided by TPs and FNs, represents that percentage of true events that were correctly predicted.

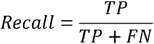

Finally, the F1-score, calculated as the harmonic mean of Precision and Recall, represents the network performance, penalizing imbalanced models.

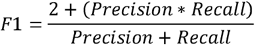

To ease subsequent evaluation of ML models for SWR analysis we provide open-access to codes for retraining strategies: https://github.com/PridaLab/rippl-AI

### Parameter fitting

Different combinations of parameters and hyper-parameters were tested for each architecture during the training phase (1944 for XGBoost, 72 for SVM, 2160 for LSTM, 60 for 2D-CNN, 576 for 1D-CNN).

Two parameters were shared across all architectures: the number of channels and the number of timestamps in the analysis window (referred as the window size). These parameters define the dimensionality of the input data (# timesteps x # channels), i.e. the number of input features.

The number of channels to be used was set at 1, 3 or 8. When 1 channel was chosen, it was that corresponding to the CA1 pyramidal layer channel, defined as the channel with most power in the ripple bandwidth (150-250 Hz). The superficial, pyramidal, and deep channels were used as 3 channels. All the channels in the shank were used for the 8-channels input configuration.

The number of timestamps defines the window size. The tested values depended on each architecture, and ranged between windows of 0.8 to 51.2 milliseconds. The rest of the parameters were specific for each architecture (see below).

The F1-score metric for the training and test set was calculated to compare the performance of the models, with the test F1 serving as a priori metric of the generalization of the models, allowing for a selection of models without performing a complete validation.

For each model, a test-F1 array was calculated with different thresholds (generally, from 0.1 to 0.9 with 0.1 increments), and the highest value for each model was used for comparison among models of the same architecture. As a result, the 50-best performing models were selected after the initial retrained test.

### Validation process

The aim of validation is to find the model that generalizes best to unseen data for each architecture. With that in mind, defining a metric that takes this into account is not a straight-forward task.

To weigh each validation session (21) independently, a F1 array was calculated for each individual session, resulting in matrix of 21 per number of threshold-values (#th). The mean of sessions gives us a #th array that quantifies the performance/generalization of the model as a function of the chosen threshold. The maximum value of this array will represent the best performance that could be achieved with this model if the threshold is correctly selected. This single value is what will be compared. Using this strategy, we narrowed down available models to the 10-best of each architecture, before selecting the best model.

### XGBoost

Based in the Gradient Boosting Decision Trees algorithm, this architecture trains a tree with a subset of samples, and then calculates its output^44^. The misclassified samples are used to train a new tree. The process is repeated until a predefined number of classifiers are trained. The final model output is the weighted combination of individual outputs.

In the training process, we worked with quantitative features (LFP values per channels) and a threshold value for a specific feature was considered in each training step. If this division correctly classifies some samples of the subset, two new nodes were generated in the next tree level, where the operation was repeated until the maximum tree depth was achieved, and a new tree with the misclassified samples is generated. The input is one dimensional (# of channels x # of timesteps) and produces a single output.

Specific parameters of XGBOOST are Maximum depth, the maximum levels for each tree, may lead to overfitting. Learning rate, which controls the influence of each individual model in the ensemble of trees. Gamma is the minimum loss reduction required to make a further partition on a leaf node, with larger values leading to conservative models. Parameter λ contributes to the regularization, with larger values preventing overfitting. Scale is used in imbalanced problems, the larger the more penalized false negatives are during training.

Trained models had a number of trainable parameters ranging from 1500 to 17900.

### SVM

Support Vector Machine is a classical classifier that searches for a hyperplane in the input dimensionality that maximizes the separation between different classes. This is only possible in lineal separation problems, so some misclassifications are allowed in real tasks. Usually, SVM performs a transformation on the original data using a kernel (linear or otherwise) that increases the data dimensionality but facilitates classification.

During training, the parameters that define the separation hyperplane are updated until the maximum number of iterations is achieved, or the rate of change in the parameters go below a threshold. The input is one-dimensional (# of channels x # of timesteps) and produces a single output.

Specific parameters of SVM are the kernel type. Using nonlinear kernels resulted in an explosive growth in training and predicting times, due to the enormous number of training data points. Only the linear kernel produced manageable times. The under-sample proportion rules out negative samples (windows without ripple) until the desired balance is achieved: 1 indicates the same number of positives and negatives.

Trained models had a number of trainable parameters ranging from 1 to 480.

### LSTM

Recurrent Neural Networks (RNNs) are a subtype of NNs especially suited to work with temporal series of data, extracting the “hidden” relations and tendencies between non-contiguous instants. Long Short-Term Memory (LSTMs) are RNNs with modifications that prevent some associated problems^46^.

During training three sets of weights and biases are updated in each LSTM unit, associated with different “gates” (Forget, input and output). To prevent overfitting, two layers of dropout (DP) and batch normalization (batchNorm) were inserted between LSTM layers. DP randomly prevents some outputs to propagate to the next layer. BatchNorm normalizes the output of the previous layer. The final layer is a dense layer that outputs the event probability. The input is two-dimensional (# of timesteps, # of channels) and produces a probability for each timestep. After each window the internal weights are reset.

Specific LSTM parameters: bidirectional if the model process the windows forwards and backwards simultaneously; # of layers is the number of LSTM layers; # of units the number of LSTM units in each layer, and # of epochs, which is the number of times the training data is used to perform training.

Trained models had a number of trainable parameters ranging from 156 to 52851.

### 2D-CNN

Convolutional Neural Networks use convolutional layers consisting of kernels (spatial filters) to extract the relevant features of an image^49^. Successive layers use this as inputs to compute general features of the image. This 2D-CNN moves the kernels along the two axes, temporal (timesteps) and spatial (channels). The first half of the architecture includes MaxPooling layers that reduce the dimensionality and prevent overfitting. A batchNorm layer follows every convolutional layer. Finally, a dense layer produces the event probability of the window.

During training, the weights and biases of every kernel are updated to minimize the loss function, with was taken as the binary cross entropy:

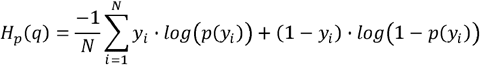

N is the number of windows in the training set, y_i_ is the label of the *i* window and p(y_i_) is the probability of ripple that the model predicts. The input is # of timesteps and # of channels; and produces a single probability for each window.

The 2D-CNN was tested with a fixed number of layers and kernel dimension. The kernel factor parameter determined the number of kernels in this structure: 32xkf (2×2), 16xkf (2×2), 8xkf (3×2), 16xkf (4×1), 16xkf (6×1) and 8xkf (8×1). In parenthesis the size of the kernels in each layer.

Trained models had a number of trainable parameters ranging from 1713 to 24513.

### 1D-CNN

This model is also a convolutional neural network, but the kernels only move along the temporal axis while processing spatial information. The number of layers and the kernel size was fixed. The tested models had 7 sets of 1D convolutional layer, batchNorm and LeakyRelu layer, followed up by a dense sigmoid activation unit. This model is similar to our previous CNN solution^31^.

During training, the weights and biases of the layers were also updated with the objective of minimizing the binary cross entropy. The input is # of timesteps and # of channels, and produces a single probability for each window.

The specific parameters for 1D-CNN included the kernel factor, which defined the number of kernels in each conv layer. The size and stride for each layer was equal and fixed. The size of the kernels in the first layer was defined as the length of the input window divided by 8. Structure: 4x*kf* (# timesteps//8 x # timesteps//8), 2x*kf* (1×1), 8x*kf* (2×2), 4×1 (1×1), 16x*kf* (2×2), 8x*kf* (1×1) and 32x*kf* (2×2). Parameters also include # of epochs, as the number of times the training data is used to perform training, and # of training batch samples, which is the number of windows that are processed before parameter updating.

Trained models had a number of trainable parameters ranging from 342 to 4253.

### Characterization of SWR features

SWR properties (ripple frequency and power) were computed using a 100 ms window around the center of the event, measured at the pyramidal channel of the raw LFP. Preferred frequency was computed first by calculating the power spectrum of the 100 ms interval using the enlarged bandpass filter 70 and 400 Hz, and then looking for the frequency of the maximum power. In order to account for the exponential power decay in higher frequencies, we subtracted a fitted exponential curve (‘fitnlm’ from MATLAB toolbox) before looking for the preferred frequency. To estimate the ripple power, the spectral contribution was computed as the sum of the power values for all frequencies lower than 100 Hz normalized by the sum of all power values for all frequencies (of note, no subtraction was applied to this power spectrum).

### Dimensionality reduction using UMAP

To classify SWR, we used topological approaches^14^. The UMAP version 0.5.1 (https://umap-learn.readthedocs.io/en/latest/) in Python 3.8.10 Anaconda was used, which is known to properly preserve local and global distances while embedding data in a lower dimensional space. In all cases, we used default values for reconstruction parameters. Algorithms were initialized randomly.

UMAP provided robust results independent on initialization.

### Prediction and retraining of non-human primate data-set

To study the generalization capabilities of the different architectures, we used data from a freely moving macaque targeting similar CA1, as completed in our mouse data (methods are described in reference^52^). Recordings were obtained with a 64-ch linear polymer probe (custom ‘deep array probe’, Diagnostic Biochips) that recorded across the CA1 layers of the anterior hippocampus (Fig. 6A) where layers were identifiable relative to the main pyramidal layer, which contains the greatest unit activity and SWP power. LFP signals were sampled at 30 kHz using a Freelynx wireless acquisition system (Neuralynx, Inc). Data corresponds to periods of immobility for a duration of almost 2 hours and 40 minutes, predominantly comprised of sleep in overnight housing. LFP intervals presenting a high level of noise across all channels was not considered for analysis.

Similar to the procedures used in mice, SWR beginning and ending times were manually tagged (ground truth). First, the best model of each architecture, already trained with the mouse data, was used to predict the output of the primate data with no retraining. For this purpose, we used recordings of different channels around the CA1 pyramidal channel, and matched to meet the laminar organization of the dorsal mouse hippocampus. Specifically, we used a CA1 radiatum channel, 720 μm from the pyramidal layer, three channels in the pyramidal layer, at +90μm, +0μm and -90μm from the pyramidal channel, and a stratum oriens channel 720μm from the pyramidal channel. The pyramidal channel was defined at the site with the maximal ripple power. We complemented these 5 recordings with 3 more interpolated signals, making a total of 8 input channels [oriens, interpolated, pyramidal, pyramidal, pyramidal, interpolated, interpolated, radiatum] using a linear interpolation script available at Github: https://github.com/PridaLab/rippl-AI/blob/main/aux_fcn.py. The applied pre-processing was the same as with the mice data: subsampling to 1250Hz and z-score normalization.

With the aim of studying the effect of retraining with completely different data, we retrained the models. Data was split in three sets (50% training, 20% test, 30% validation), and used to retrain and validate the models. For re-training, we reset all trainable parameters (internal weights) but kept all architectural hyper-parameters fixed (input number of channels, input window length, number of layers, etc…) as with the mouse data, making the re-training process much faster than the original training that required a deep hyper-parametric search (per model re-train: 2min for XGBoost, 10-30min for SVM, 3-20min for LSTM, 1-10 min for 2D-CNN and 1-15 min for 1D-CNN).

## Code and data availability

Data and codes used in this study are available. The training and test set data are available at https://figshare.com/authors/Liset_M_de_la_Prida/402282 and listed independently as follows:

M de la Prida, Liset (2021): Amigo2_2019-07-11_11-57-07. figshare. Dataset. https://doi.org/10.6084/m9.figshare.16847521.v2

M de la Prida, Liset (2021): Som2_2019-07-24_12-01-49. figshare. Dataset. https://doi.org/10.6084/m9.figshare.16856137.v2

M de la Prida, Liset (2021): Dlx1_2021-02-12_12-46-54. figshare. Dataset. https://doi.org/10.6084/m9.figshare.14959449.v4

M de la Prida, Liset (2021): Thy7_2020-11-11_16-05-00. figshare. Dataset. https://doi.org/10.6084/m9.figshare.14960085.v1

Codes for some of the best trained models of all architectures are available in an open-source repository https://github.com/PridaLab/rippl-AI and documented in open-source notebooks for model retraining https://github.com/PridaLab/rippl-AI/blob/main/examples_retraining.ipynb and for SWR detection https://github.com/PridaLab/rippl-AI/blob/main/examples_detection.ipynb

## Acknowledgements

This work was supported by the Fundación La Caixa (LCF/PR/HR21/52410030) to LMP and by the Whitehall Foundation and BRAIN Initiative NINDS (R01NS127128) to KLH. We thank the Spanish Society of Neuroscience (SENC) and the Universidad Autónoma de Madrid Doctorate for partially supporting the hackathon. ANO was supported by PhD fellowships from the Spanish Ministry of Education (FPU17/03268). AR was supported by a JAE-Intro Fellowship of the AI-HUB CSIC program (JAE Intro AI HUB21) and by the CSIC Interdisciplinary Thematic Platform Neuro-Aging (PTI+Neuro-Aging). We thank all participants of the hackathon. Thanks to Rodrigo Amaducci, Enrique R Sebastian, Daniel García-Rincón and Adrián Gollerizo for co-organizing the hackathon.

